# Cep97 Is Required For Centriole Structural Integrity And Cilia Formation In *Drosophila*

**DOI:** 10.1101/740217

**Authors:** Jeroen Dobbelaere, Marketa Schmidt-Cernohorska, Martina Huranova, Dea Slade, Alexander Dammermann

**Affiliations:** Max Perutz Labs, University of Vienna, Vienna Biocenter (VBC), Dr Bohr-Gasse 9, A-1030 Vienna, Austria; Laboratory of Adaptive Immunity, Institute of Molecular Genetics of the Czech Academy of Sciences, Videnska 1083, Prague 14220, Czech Republic

## Abstract

Centrioles are highly elaborate microtubule-based structures responsible for the formation of centrosomes and cilia. Despite considerable variation across species and tissues, within any given tissue their size is essentially constant [1, 2]. While the diameter of the centriole cylinder is set by the dimensions of the inner scaffolding structure of the cartwheel [3], how centriole length is set so precisely and stably maintained over many cell divisions is not well understood. Cep97 and CP110 are conserved proteins that localize to the distal end of centrioles and have been reported to limit centriole elongation in vertebrates [4, 5]. Here, we examine Cep97 function in *Drosophila melanogaster*. We show that Cep97 is essential for formation of full-length centrioles in multiple tissues of the fly. We further identify the microtubule deacetylase Sirt2 as a Cep97 proximity interactor. Deletion of Sirt2 likewise affects centriole size. Interestingly, so does deletion of the acetylase Atat1, indicating that loss of stabilizing acetyl marks impairs centriole integrity. Cep97 and CP110 were originally identified as inhibitors of cilia formation in vertebrate cultured cells [6] and loss of CP110 is a widely used marker of basal body maturation. In contrast, in *Drosophila* Cep97 is only transiently removed from basal bodies and loss of Cep97 strongly impairs ciliogenesis. Collectively, our results support a model whereby Cep97 functions as part of a protective cap that acts together with the microtubule acetylation machinery to maintain centriole stability, essential for proper function in cilium biogenesis.

## RESULTS AND DISCUSSION

Centrioles generally assemble adjacent to pre-existing parental centrioles in a series of steps which have been extensively studied in a range of experimental models. First, PLK4/Sak/ZYG-1 is recruited to the vicinity of the mother centriole where it concentrates into a single focus that marks the site of daughter centriole assembly. PLK4 then recruits and phosphorylates STIL/Ana2/SAS-5 as well as SAS-6, which oligomerizes to form the hub-and-spoke structure of the cartwheel. Finally, CPAP/SAS-4 along with *γ*-tubulin directs the assembly of the microtubule-based centriole wall, aided in some species by CEP135/Bld10 [7]. Overexpression of CPAP as well as its interacting proteins CEP120 and SPICE1 is known to result in over-elongation of centrioles [4, 5, 8–10] and recent studies have demonstrated that CPAP plays a key role in imparting slow, processive growth on centriolar microtubules [11, 12]. The distal (plus) ends of centriolar microtubules are bound by a complex of Cep97 and CP110 [6, 13, 14]. In vertebrates, this complex is thought to counteract CPAP activity, as loss of either component results in over-elongated centrioles, while overexpression of CP110 can suppress the effect of excess CPAP [4, 5]. Cep97 and CP110 have been shown to interact with the depolymerizing kinesin KIF24, which specifically acts on centriolar, but not cytoplasmic microtubules to limit centriole size [15]. An interaction between CP110 and another depolymerizing kinesin, Klp10A, was reported in *Drosophila* and loss or depletion of Klp10A results in a dramatic extension of centriole size, an effect suppressed by overexpression of CP110 [16, 17]. Surprisingly, however, loss of CP110 in the fly has only minor effects on centriole elongation and, indeed, results in shortening of centrioles in Dmel cells, hinting at context-dependent differences [16, 17]. Further, while Cep97 and CP110 are generally thought to function together, it has been noted that hypomorphic *Cep97* mutants display stronger phenotypes than a *CP110* null mutant, suggesting a role for Cep97 independent of CP110 [18]. We therefore sought to carry out a comprehensive analysis of Cep97 in the fly, using newly developed tools to monitor protein localization and function.

### Cep97 stably caps mature centrioles

Both in vertebrates and *Drosophila* Cep97 has so far exclusively been studied by immunofluorescence microscopy in fixed cells or using fluorescent fusions expressed under a heterologous promoter. To examine the localization and *in vivo* dynamics of Cep97 without the potential complications of overexpression, we generated a GFP fusion under the control of endogenous regulatory sequences by recombineering and φC31 integrase-mediated insertion at a defined chromosomal locus. This fusion was found to be fully functional in rescuing the phenotypes of Cep97 deletion (see below) and expressed at levels similar to the endogenous protein (Fig. 2A). Cep97 was found to localize to centrioles throughout the animal (Fig. 1A). A detailed examination of the spatial distribution on the giant centrioles in spermatocytes by STED microscopy found Cep97 concentrated in the distal lumen, with a full width at half maximum (FWHM) of 100±38nm (N=11) (Fig. 1B), consistent with that of *Drosophila* CP110 [17, 19], but considerably narrower than the distribution reported for both proteins in human retinal pigment epithelial (RPE) cells (172±35nm and 196±24nm, respectively, approximately the diameter of the centriole wall [14]). The reasons for this discrepancy are not immediately clear. However, we could confirm these findings by expansion microscopy on RPE cells in our own hands (172±22nm and 190±24nm, respectively, see Fig. S1A, D).

**Figure 1:**
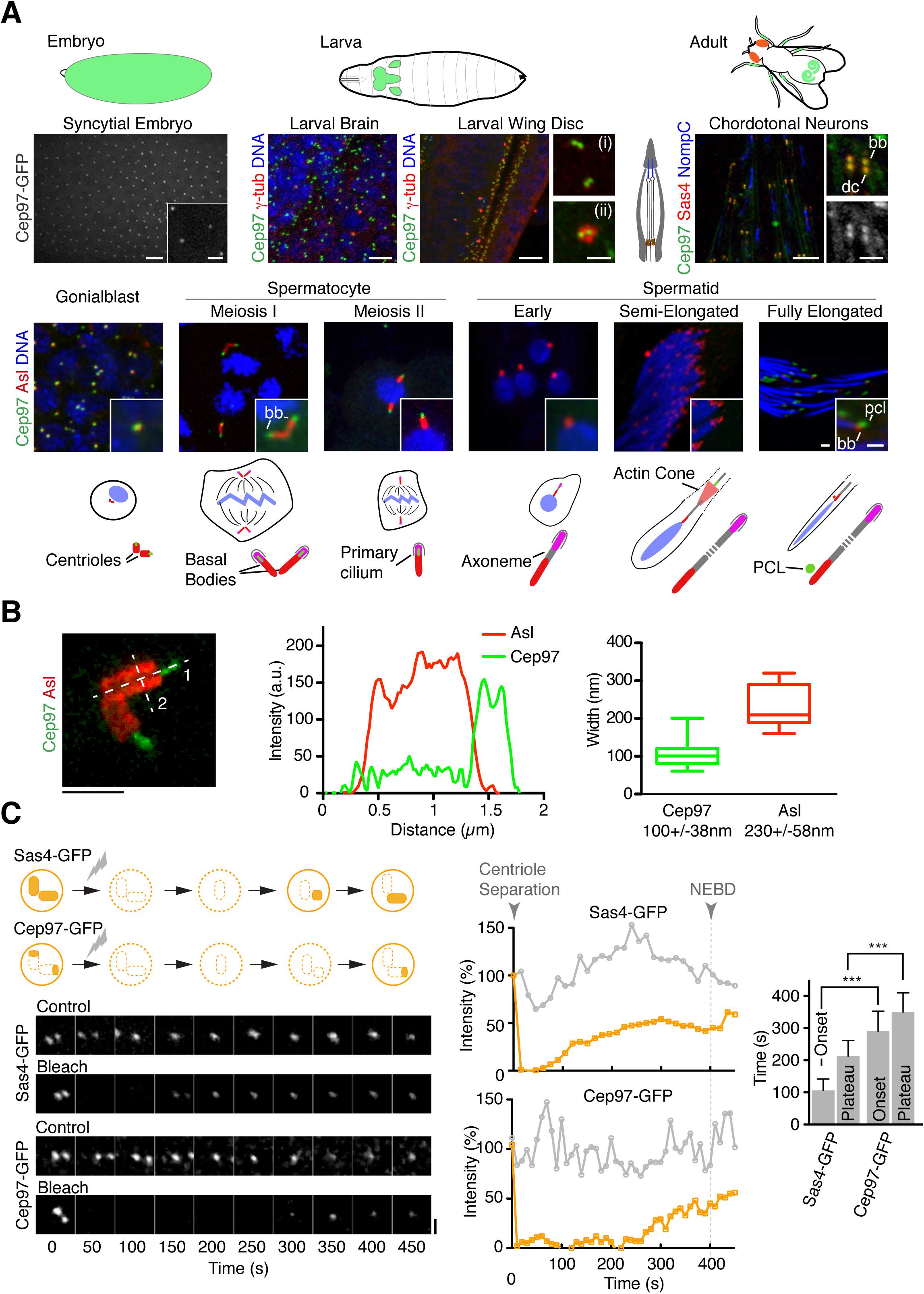
Cep97 stably localizes to mature centrioles and basal bodies in *Drosophila*. (**A**) Schematics of all tissues used to determine Cep97 localization and function (centrioles in syncytial-stage embryo, larval brain and wing disc, basal bodies in adult leg chordotonal organs and testes). Immunofluorescence micrographs show Cep97 co-localizing with centriolar/centrosomal markers (Asl, Sas4, *γ*-tubulin) in all tissues examined, including at the basal bodies of chordotonal neurons (bb, indicated; dc, daughter centriole) and primary spermatocytes and the procentriole-like structure (PCL) of mature sperm, though not the sperm basal body. Signal unchanged between interphase (i) and mitosis (ii). NompC used to visualize ciliary membrane in chordotonal neurons. Insets acquired in airyscan mode. Scale bars are 5µm, 1µm insets. (**B**) Cep97 localizes to the distal lumen of centrioles. Immunofluorescence micrograph of spermatocyte centriole pair acquired by stimulated emission depletion (STED) microscopy along with linescan of the fluorescence signal distribution along the proximal-distal axis (1, middle) and measurement of the lateral distribution for Cep97 and the centriolar wall marker Asl (2, right). Lateral distribution measured as full width at half maximum, Cep97, N=11, peak-peak distance, Asl, N=12. Scale bar is 1µm. Box and whisker plot with whiskers representing minimum and maximum values. (**C**) Fluorescence recovery after photobleaching (FRAP) in syncytial-stage embryos shows stable incorporation of Cep97 with delayed recruitment relative to Sas4. Schematic of FRAP assay, representative images and recovery curves for Sas4-GFP and Cep97-GFP. Photobleaching was performed just after centriole separation and the time of initial signal recovery (i.e. new recruitment) and the time at which a new plateau is reached recorded. Plots are for single FRAP experiment, displaying average of two bleached centrioles (orange) and four control centrioles (grey). N=12 Sas4-GFP, 8 Cep97-GFP. Scale bar is 1µm. Error bars are SD. *** t-test, P<0.001.

**Figure 2:**
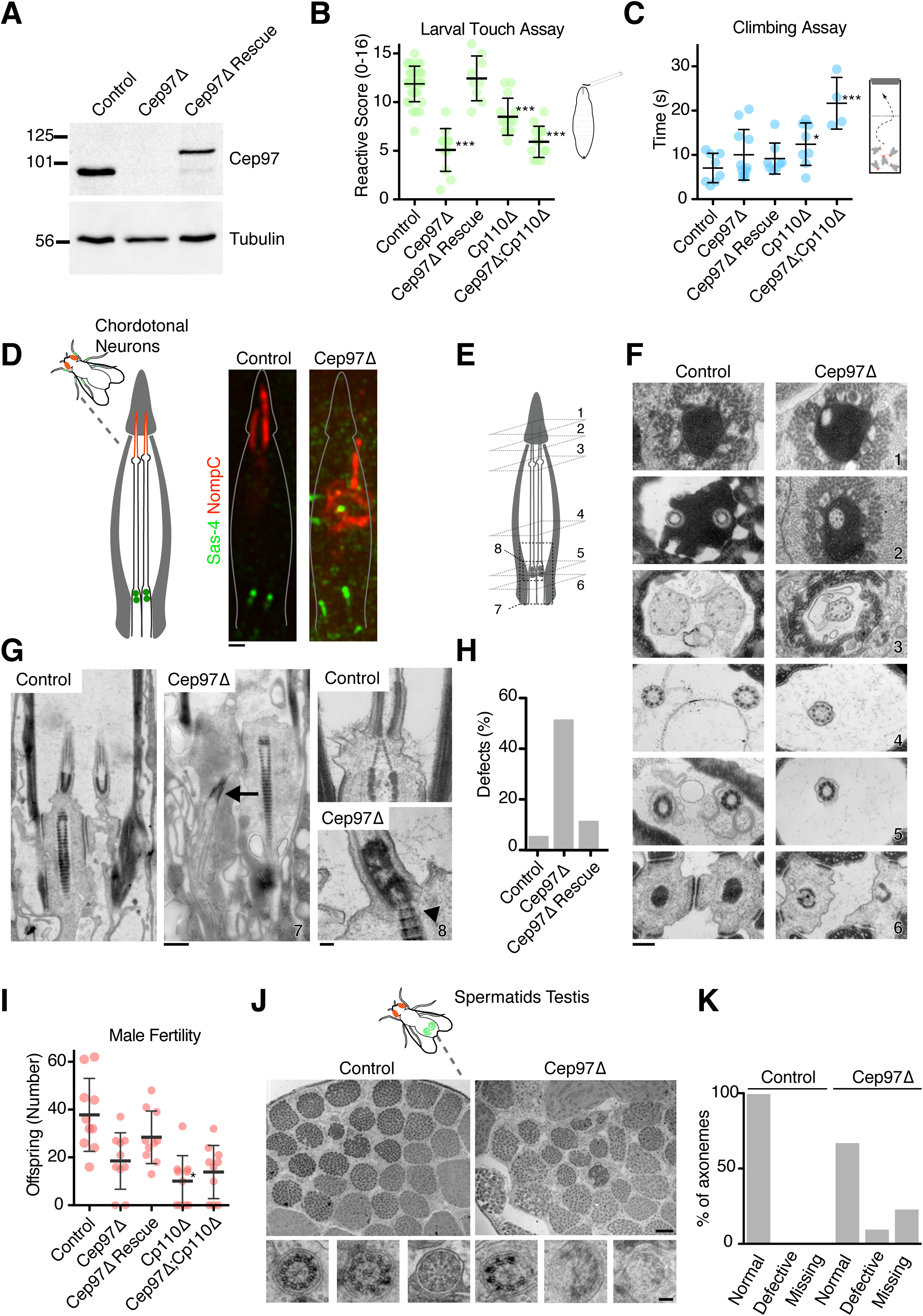
Cep97 deletion mutants display ciliary defects. (**A**) Immunoblotting confirms loss of Cep97 in *Cep97Δ* animals and expression of endogenous promoter Cep97-GFP transgene to levels similar to controls. (**B, C**) Behavioral assays in larvae (touch assay, **B**) and adult flies (climbing/bang assay, **C**) show loss of coordination in *Cep97* and *CP110* mutants, rescued for *Cep97* by expression of Cep97-GFP. Error bars are SD. * t-test, P<0.05, *** P<0.001. (**D**) Schematic and immmunofluorescence micrographs of scolopidia in chordotonal organ of the fly. Sas-4 and NompC used to visualize basal body and ciliary tip, respectively. Each scolopidium contains two cilia with their ends embedded in the cap cell. *Cep97* mutants display severe disorganization of ciliary morphology. Scale bar is 1µm. (**E**-**G**) Cross (**F**) and longitudinal sections (**G**) through control and *Cep97* mutant scolopidia analyzed by transmission electron microscopy (TEM). Positions are indicated by numbers in schematic in (**E**). Scale bars are 200nm (**F**, panels 1-6), 500nm (**G**, 7) and 100nm (**G**, 8). Cross sections reveal missing cilium at all levels, though remaining cilium appears morphologically normal. Longitudinal sections reveal mispositioned basal bodies (arrow) as well as rootlets emanating from docked basal body rather than daughter centriole as in controls (arrowhead). See also Fig. S3A. (**H**) Quantitation of ciliary defects revealed by ultrastructural analysis in Cep97 mutants and GFP rescue animals. N=90 control, 118 *Cep97Δ*, 95 *Cep97Δ* Rescue. (**I**) *Cep97* and *CP110* mutants display reduced male fertility, assessed as number of offspring per single male. Error bars are SD. * t-test, P<0.05. (**J, K**) TEM analysis of sperm axonemes in control and *Cep97* mutant testes. Cross sectional views (**J**) reveal missing axonemes (each cyst normally contains 64 sperm developing synchronously), as well as fragmented axonemes in Cep97 mutants. Scale bars are 5µm, 100nm insets. Defects quantitated in (**K**). N=16 control, 11 *Cep97Δ*.

We next examined the recruitment dynamics of Cep97 in syncytial embryos, which rapidly cycle between S- and M-phases. New centrioles assemble in each S phase and separate at the end of the subsequent mitosis. While mother and daughter centrioles prior to this remain in close proximity and cannot be resolved by standard confocal microscopy, previous work has shown that the timing of recruitment and turnover of centriolar proteins can be accurately assessed by fluorescence recovery after photobleaching (FRAP) [20]. When applied to Sas4, a component stably incorporated into the centriolar microtubule wall [21], this assay shows recovery of centriolar signal beginning ∼100s after mitotic exit and reaching 50% of the original signal (Fig. 1C), reflecting recruitment to newly forming centrioles (Sas4-GFP on the original parental centriole remaining photobleached and accounting for the missing 50%). In contrast, Cep97 signal does not become begin to recover until ∼300s, at which point Sas4 signal has reached a plateau and daughter centrioles have reached their mature length ([13], Fig. 1C). The failure to reach the original pre-bleach signal intensity further indicates that centriolar Cep97, like Sas4 and Sas6 [13, 21, 22], does not exhibit cytoplasmic exchange and therefore forms a stable cap at the distal end of fully grown centrioles. It should be noted that these results conflict with those reported by [13], which found *Drosophila* Cep97 dynamically associated with the growing end of centrioles, which we believe reflects an excess of Cep97 due to the use of a strong ubiquitin promoter to drive GFP expression [23].

### Cep97 is not universally stripped from the ciliary base

Removal of CP110 from the maturing basal body is a hallmark event in vertebrate ciliogenesis, occurring after distal appendage-mediated ciliary vesicle docking and requiring the activity of tau tubulin kinase (TTBK2) [24, 25]. This removal appears to trigger transition zone assembly and extension of the ciliary axoneme (reviewed in [26]). While most work has focused on loss of CP110, Cep97 appears to follow the same pattern [5] (see also Fig. S1B, C). We first examined Cep97 localization in spermatogenesis (Fig. 1A). Spermatogenesis in *Drosophila* can be divided into three stages: a mitotic stage in which male germ cells divide asymmetrically to generate goniablasts, which in turn divide four more times to generate a cyst of 16 primary spermatocytes; a meiotic stage in which these primary spermatocytes divide to produce 64 haploid spermatids; and finally, a differentiation stage in which spermatids mature into spermatozoa [27]. Centrioles elongate considerably during spermatogenesis, doubling in size between the early spermatocyte stage and meiosis I. They also form a short primary cilium, which is partially resorbed and internalized at the onset of meiosis [28]. Upon meiotic exit at the spermatid stage, further remodeling eventually results in the formation of the sperm flagellum. This process involves separation of the transition zone from the basal body [29]. Additionally, a procentriole-like structure transiently forms adjacent to the maturing basal body [30]. In contrast to CP110, removed at the onset of centriole elongation in early spermatocytes (Tates stage 5, [17]), Cep97 persists at the distal end of centrioles throughout early spermatogenesis up to and including meiosis (Fig. 1A). Removal of Cep97 is therefore clearly not required for initial basal body elongation or formation of the primary cilium in primary spermatocytes. Cep97 is, however, lost as basal bodies are remodeled to give rise to the sperm flagellum (Tates stage 12, Fig. 1A). While this coincides with migration of the transition zone, we found no colocalization with the transition zone component Cep290, a reported CP110 interactor in vertebrates [31] during migration of the transition zone (Fig. S1E). Late in spermatocyte maturation Cep97 was, however, observed along the ciliary axoneme ahead of the traveling actin cones that mediate sperm individualization (Fig. S1F). Recruitment to the elongating axoneme by end-binding proteins including the Cep97 interactor CEP104 has been proposed as a means for removal of Cep97/CP110 [32], but has hitherto not been observed in unperturbed cells. Finally, as in the case of primary spermatocytes, Cep97 was still localized to the basal body of mechanosensory primary cilia in leg chordotonal neurons (Fig. 1A). Collectively, these results indicate that Cep97 need not be removed for centriole elongation or primary ciliogenesis to occur. Although this clearly goes against the dogma in the field, it is not entirely unprecedented since CP110 has been reported to be still present at the mature basal bodies of *Xenopus* multiciliated cells [33]. It is also potentially significant given the requirement of Cep97 for proper ciliogenesis (see below).

### Cep97 mutant flies are viable but uncoordinated

Previous work on *Drosophila* Cep97 employed a transposable element insertion in the first intron, which was shown to behave as a hypomorph [18]. To obtain a true null mutant, we used the ends-out method [34] to replace the entire coding sequence of Cep97 with the White marker gene (Fig. S2A). We obtained several independent lines, all of which lost the Cep97 coding region according to PCR and Southern blot analysis (Fig. S2B, C and not shown). In homozygote animals, Cep97 mRNA and protein was undetectable by RT-PCR and Western blotting (Fig 2A and S2D). As was shown to be the case for *CP110* mutants [17], *Cep97* null flies were found to be viable and fertile, with no obvious morphological defects (Fig. S2E). However, mutants displayed behavioral defects characteristic of impaired mechanosensory function. Thus, touch sensation in larvae [35] was found to be strongly reduced (Fig. 2B). Similarly, adult flies displayed defects in negative geotaxis as assessed in the climbing assay (Fig. 2C), which monitors recovery from mechanical agitation [36]. Mutants also displayed reduced male fertility (Fig. 2I). Defects were rescued by introduction of Cep97-GFP, confirming specificity of the mutant phenotype and functionality of our GFP transgene.

### Cep97 mutants are cilia defective

Mechanosensory neurons in the fly are ciliated, raising the possibility that behavioral defects stem from defects in ciliary architecture. We therefore examined the sensory organs responsible for proprioception in *Drosophila*, the chordotonal organs, in the femur of the leg. Chordotonal organs are made of multiple scolopidia, each of which contains a pair of ciliated nerve endings ensheathed by a scolopale glia cell and attached with their ciliary tips to the cuticle via a cap cell [37]. Antibody staining for the centriolar protein Sas-4 and the TRP channel protein NompC, which localizes to the distal end of mechanosensory cilia [38], in *Cep97* mutants showed that centrioles/basal bodies are still present but cilia appear to be largely missing (Fig. 2D). To analyze these defects in more detail, we examined ciliary ultrastructure in *Cep97* mutant flies and wild-type controls (Fig. 2E-H). Cross-sectional views of the chordotonal organs of wild-type flies always showed a pair of cilia, positioned side by side with their tips embedded in the cap cell. In contrast, one or both cilia were frequently missing in *Cep97* mutants (Fig. 2F, H). In cases where both cilia were still present, they frequently appeared to be laterally displaced with landmark features such as the ciliary dilation found in different sections, suggesting defects in basal body positioning. Longitudinal sections support this idea, with what appear to be maloriented basal bodies found in certain sections (Fig. 2G, S3). Close-up views of individual structures revealed no marked defects of the axoneme, transition zone or rootlets (Fig. 2F). However, basal bodies frequently appeared abnormal, with mother and daughter centrioles not always clearly distinct and instances of rootlets apparently emanating from the mother centriole, not the daughter as usually the case in *Drosophila*. Other centriole pairs did not appear to be associated with ciliary structures (Fig. 2G, S3). Collectively, these results suggest Cep97 is required for the proper formation of cilia, with basal body function apparently impaired in *Cep97* mutants.

**Figure 3:**
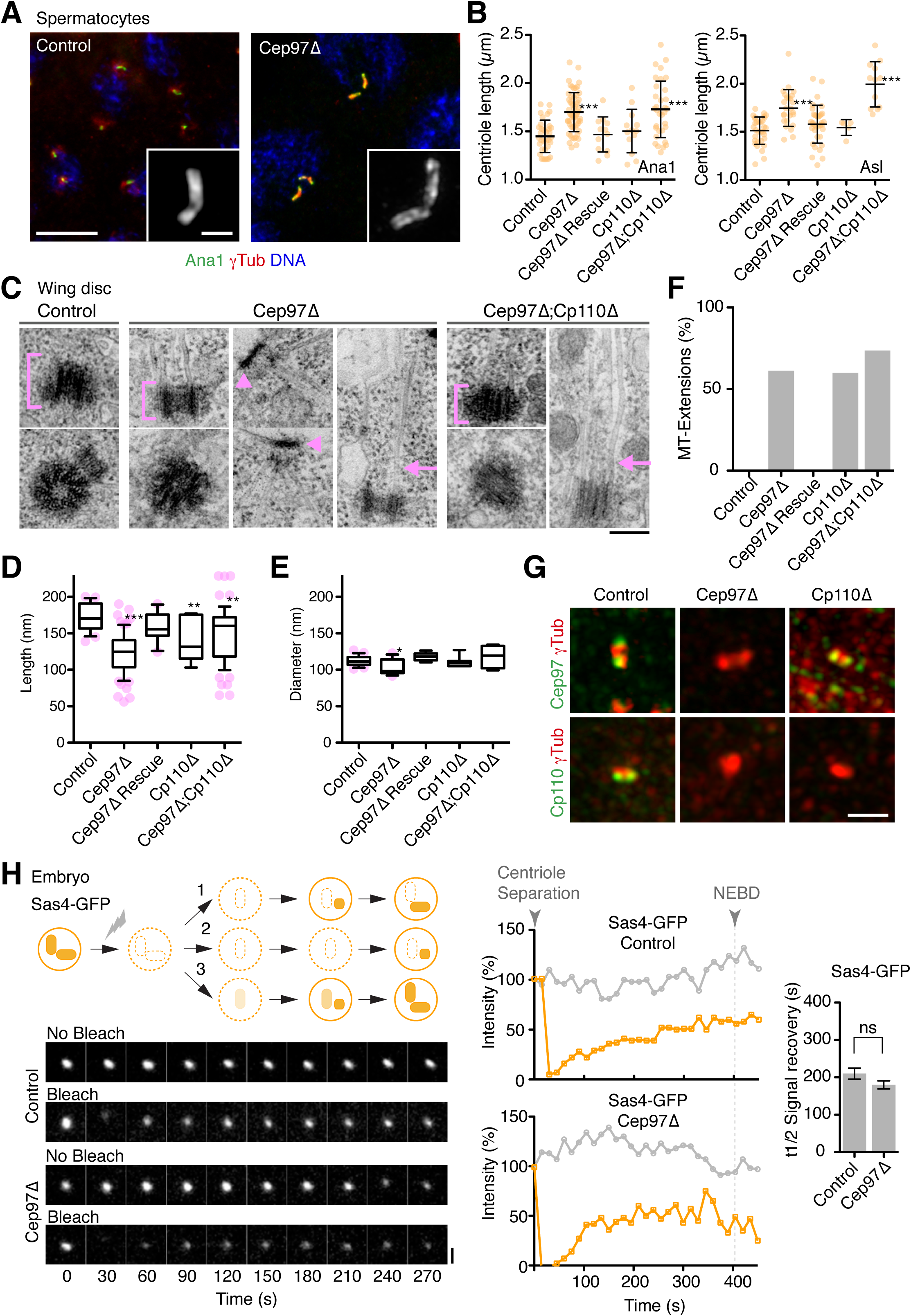
Cep97 is required for centriole stability. (**A**) Immunofluorescence micrographs of control and *Cep97* mutant spermatocytes stained as indicated. Centrioles appear visibly longer than in controls as assessed using the centriolar marker Ana1. Insets acquired in airyscan mode. Scale bars are 10µm, 1µm insets. (**B**) Quantitation of spermatocyte centriole lengths on images as in (**A**) using Ana1 or Asl as centriolar markers. Note that significant centriole elongation is observed only in *Cep97* mutants, not mutants of *CP110*. Error bars are SD. *** t-test, P<0.001. (**C**-**F**) Transmission electron micrographs of centrioles in control, *Cep97* mutant and *Cep97;CP110* double mutant wing discs. *Cep97* and *CP110* mutant centrioles are more variable in length but generally shorter than in controls, while diameter is not affected (quantified in **D, E**; N=44/52 control (**D**/**E)**, 71/15 *Cep97Δ*, 17/8 *Cep97Δ* Rescue, 15/7 *CP110Δ*, 69/9 *Cep97Δ;CP110Δ*; box and whisker plot with whiskers representing 10th and 90th percentiles; single dots represent outliers; ** t-test, P<0.01, *** P<0.001). Individual microtubules are seen extending from the centriole barrel in both mutants (arrows; quantified in **F**, N=39 control, 65 *Cep97Δ*, 18 *Cep97Δ* Rescue, 35 *CP110Δ*, 56 *Cep97Δ;CP110Δ*). *Cep97* mutants further display disc-like structures acting as MTOCs which may be severely truncated centrioles (arrowheads). Scale bar is 100nm. (**G**) Centriolar localization interdependencies between Cep97 and CP110. Immunofluorescence micrographs of Cep97 and CP110 in control and mutant spermatocytes as indicated. *γ*-tubulin is used as a countermarker. Cep97 is required for CP110 localization but not vice versa. Images acquired in airyscan mode. Scale bar is 1µm. (**H**) Fluorescence recovery after photobleaching (FRAP) performed on Sas4-GFP-expressing syncytial-stage embryos as in Fig. 1C. Sas4 is normally stably incorporated during centriole assembly (1). Lack of Cep97 could theoretically manifest itself in a failure to recruit Sas4 with normal kinetics (2) or failure to stably incorporate Sas4 (3). Actual results reveal no change in Sas4 stable incorporation or recruitment kinetics (N=12 Control, 16 *Cep97Δ*). Scale bar is 1µm. Error bars are SD.

Since male fertility was also affected, we wondered whether these defects also extend to the sperm flagellum. Examining the testes of wild type flies by thin section transmission electron microscopy, we found multiple cysts of 64 spermatids at different stages of development. In *Cep97* mutants, early spermatid cysts displayed a similar number of spermatids (12/12 cysts with 64 cells), indicating proper completion of meiosis (Fig. 2J and data not shown). However, even at early stages abnormal spermatids could be observed in which axonemes were incomplete or entirely missing. These defects became more pronounced in mature spermatids, consistent with impaired axoneme elongation and/or stability (Fig. 2J, K). Thus, Cep97 is also required for proper ciliogenesis in sperm. However, a sizeable number of structurally normal axonemes was still observed and live microscopy showed the presence of motile sperm in the seminal vesicle (not shown), consistent with the residual fertility of *Cep97* mutant males.

While both Cep97 and CP110 were originally described as inhibitors of ciliogenesis, it has recently been reported that CP110 also has a positive role in promoting ciliogenesis, with impaired docking of basal bodies following CP110 depletion/mutation in vertebrates [33, 39]. Our results are consistent with this idea and suggest Cep97 is likewise required for proper basal body function.

### Cep97 is required for assembly of proper length centrioles

Defective basal body function likely stems from structural defects in the underlying centriole template. Centriole numbers were found to be unchanged in mutant embryos, suggesting centriole duplication is not appreciably affected (not shown). PCM recruitment and mitotic progression were also unaffected. Centrioles likewise appeared superficially normal in the early stages of spermatogenesis. However, as centrioles elongate in primary spermatocytes, Cep97 mutant centrioles grew to a length of 1.70µm as measured by immunofluorescence using the centriolar markers Ana1 and Asl, significantly longer than the 1.45µm found in wild-type controls (Fig. 3A, B). This marked size difference was maintained for the remainder of spermatogenesis and still evident for mature basal bodies (not shown). Interestingly, centrioles in wing discs, a non-ciliated somatic tissue, behaved very differently. Here, electron microscopy reveals dimensions of 172.4 ± 3.9nm (length) by 111.9 ± 1.3nm (width) for wild-type centrioles (Fig. 3C-E). In contrast to what we found in sperm, Cep97 mutant centrioles were markedly shorter (123 ± 3.3nm), a phenotype rescued by expression of Cep97-GFP. This 30% reduction in length is likely an underestimate as Cep97 mutant wing discs further displayed numerous disk-like structures not found in controls that may be highly truncated centrioles (see Fig. 3C). However, since these structures could not be unambiguously identified as centrioles, they were excluded from our analysis. Interestingly, both truncated centrioles and disk-like structures displayed prominent extensions of individual microtubules emanating from one end (Fig. 3C, F). A similar phenomenon had been previously reported for CP110 [17]. However, *CP110* mutants otherwise displayed much less severe phenotypes, with centriole length only slightly different from wild-type in both spermatocytes and wing-discs ([17] and Fig. 3B, D). Further, *Cep97;CP110* double mutants displayed a phenotype essentially indistinguishable from that of *Cep97* single mutants. Cep97 is generally thought of as purely a recruitment factor for CP110 [6]. Cep97 indeed does recruit CP110 to centrioles also in the fly (Fig. 3G). However, our results clearly indicate that Cep97 has other functions in centriole length control besides CP110 recruitment. Further, both Cep97 and CP110 are generally thought to act to limit centriole elongation by opposing the activity of proteins such as CPAP [4, 5]. This has also been reported for CP110 in *Drosophila* [17]. Yet, the consequences of Cep97 loss are clearly more differentiated, with both abnormal elongation (spermatocytes) and shrinkage (wing discs) occurring depending of cellular context, with a higher degree of variability observed in both cases. As previously proposed by the Glover lab [16], we therefore see the primary role of Cep97 and CP110 not as limiting centriole elongation but rather acting as a protective cap limiting further centriole growth or potentially shrinkage once the proper length has been reached.

### Cep97 interacts with the microtubule acetylation machinery to stabilize centrioles

How might Cep97 and CP110 function in capping centrioles? Given the stable association of Cep97 with the distal end of centrioles, a direct effect on centriolar microtubule turnover cannot be excluded. However, an examination of the dynamics of Sas4, a component of the centriolar wall, by fluorescence recovery after photobleaching in syncytial stage embryos as described above, reveals no obvious difference in the extent of recruitment in Cep97 mutants compared to controls (Fig. 3H). Furthermore, Sas4 continues to be stably incorporated, with no detectable cytoplasmic exchange on fully mature centrioles. This is in contrast to the consequences of depletion of *γ*-tubulin or tubulin itself, which results in Sas4 remaining partially or fully dynamic [21]. Therefore, the early stages of centriole assembly up to and including centriolar microtubule wall assembly appear to be largely unaffected by loss of Cep97.

What, then might explain the stabilizing influence of Cep97? A hint came from an unexpected source, an effort to identify interactors of the lysine deacetylase Sirt2 using a combined tandem affinity purification (TAP) and proximity biotinylation (BioID) approach [40]. Sirt2 is known to shuttle between nucleus and cytoplasm, where it colocalizes with the microtubule cytoskeleton and along with another enzyme Hdac6 acts to deacetylate cytoplasmic microtubules [41, 42]. Cep97 was found as one of the top TAP/BioID interactors of Sirt2 in cytoplasmic fractions of human cell extracts [40]. Reciprocal pulldowns of FLAG-tagged Cep97 from HEK293T cells reliably recovered co-transfected HA-Sirt2, confirming this interaction (Fig. 4A). Acetylation of lysine 40 on *α*-tubulin mediated by Atat1 is a post-translational modification associated with stable microtubules, including those at centrioles and cilia, although the precise functional significance of this modification remains poorly understood [43, 44]. RNAi-mediated depletion of Atat1 has been reported to delay ciliogenesis in serum-starved vertebrate cultured cells [45], while Hdac6 and Sirt2 are thought to function in cilia resorption upon resumption of the cell cycle [46, 47]. To study the potential role of Atat1, Sirt2 and Hdac6 in the fly, we obtained putative loss of function mutants in their respective *Drosophila* homolog. As previously observed for *Cep97*, we found all three mutants to be viable, with no obvious morphological defects (Fig. S4A, B). However, all three also displayed signs of uncoordination, with impaired response to touch in larvae and defects in negative geotaxis in adults, as well as reduced male fertility (Fig. S4C-E). Ultrastructural analysis of mutant sperm revealed a similar range of axonemal defects as for *Cep97*, albeit at lower frequencies (Figs. 4B, C, S4F). Double mutants of *Cep97* with *Sirt2* and *Hdac6* did not result in any synergistic phenotypes, either at the whole animal behavior or cilia morphology level (Figs. 4B, C, S4C-F). However, co-deletion of *Cep97* and *Atat1* was synthetic lethal (Fig. S4B). Double mutants of *Cep97* with a weaker transposon insertion mutant of *Atat1* (*Atat1-P*) were viable, but displayed strongly enhanced phenotypes, including severe uncoordination and fully penetrant male infertility (Fig. S4C-E). Consistent with this, examination of *Cep97;Atat1* double mutant sperm by transmission electron microscopy revealed almost complete disruption of axonemal architecture (Figs. 4B, C, S4F). Thus, Cep97 and components of the microtubule acetylation machinery share similar ciliary phenotypes in the fly, although the observed synergism between Cep97 and the acetylase Atat1 also suggest independent functions of each protein.

**Figure 4:**
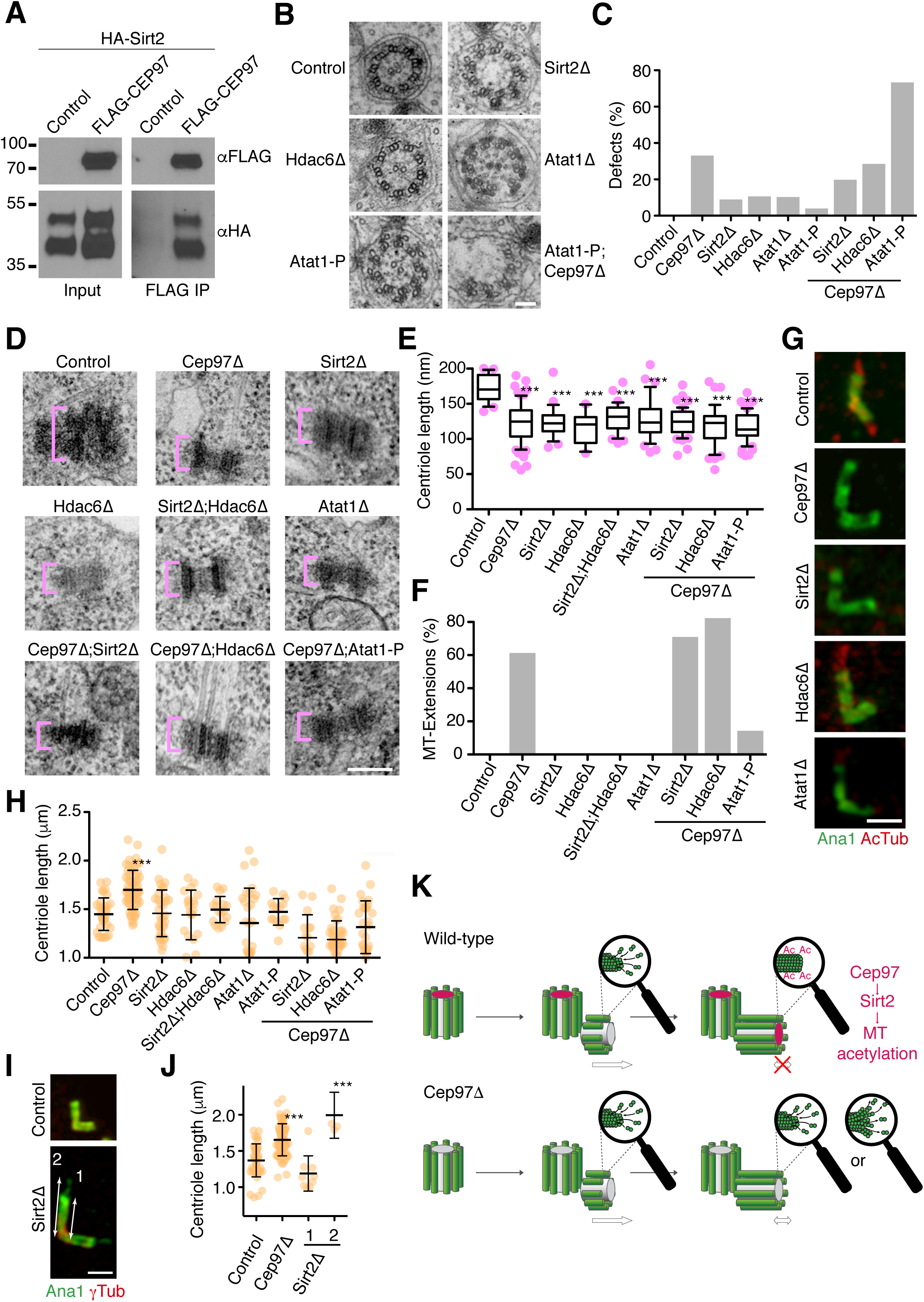
Cep97 contributes to centriolar microtubule acetylation via Sirt2. (**A**) Pulldowns of FLAG-Cep97 from HEK293T cells efficiently co-immunoprecipitate HA-Sirt2. Controls are FLAG-empty vector. (**B, C**) TEM analysis of sperm axonemes mutant for the acetylase *Atat1* and the deacetylases *Sirt2* and *Hdac6* reveal defects similar to *Cep97* (compare Fig. 2J). Cross sectional views (**B**) reveal fragmented axonemes, albeit at lower frequencies than in *Cep97* mutants (**C**). Defects are compounded in *Cep97;Atat1* double mutants (note that this is for the hypomorphic *Atat1-P* allele as *Cep97;Atat1Δ* is synthetic lethal). Scale bar is 100nm. N=22 *Sirt2Δ*, 17 *Hdac6Δ*, 18 *Atat1Δ*, 26 *Atat1-P*, 19 *Cep97Δ;Sirt2Δ*, 16 *Cep97Δ;Hdac6Δ*, 17 *Cep97Δ;Atat1-P*. Control and *Cep97Δ* data from Fig. 2K shown for comparison. See also Fig. S4F. (**D**) Transmission electron micrographs of centrioles in acetylation mutant wing discs. Mutant centrioles are more variable in length but generally shorter than in controls (quantified in **E**; N= 25 *Sirt2Δ*, 12 *Hdac6Δ*, 35 *Sirt2Δ;Hdac6Δ*, 33 *Atat1Δ*, 50 *Cep97Δ;Sirt2Δ*, 43 *Cep97Δ;Hdac6Δ*, 61 *Cep97Δ;Atat1-P*; Control and *Cep97Δ* data from Fig. 3D shown for comparison; box and whisker plot with whiskers representing 10th and 90th percentiles; single dots represent outliers; *** t-test, P<0.001). Scale bar is 100nm. (**F**) Microtubule extensions found for *Cep97* and *CP110* (Fig 3C) are not seen in *Atat1*, *Sirt2* or *Hdac6* mutants and are suppressed in *Cep97;Atat1* double mutants. N= 30 *Sirt2Δ*, 19 *Hdac6Δ*, 24 *Sirt2Δ;Hdac6Δ*, 35 *Atat1Δ*, 54 *Cep97Δ;Sirt2Δ*, 49 *Cep97Δ;Hdac6Δ*, 56 *Cep97Δ;Atat1-P*. Control and *Cep97Δ* data from Fig. 3F shown for comparison. (**G**) Immunofluorescence micrographs of *Cep97*, *Sirt2*, *Hdac6* and *Atat1* mutant spermatocyte centrioles stained for acetylated tubulin and Ana1. Microtubule acetylation is strongly reduced in all four mutants. Images acquired in airyscan mode. Scale bar is 1µm. (**H**) Centriole length in acetylation mutant spermatocytes measured on images as in (**G**). Centriole length is more variable but on average not significantly different from controls. Co-mutation of acetylation machinery suppresses centriole elongation associated with *Cep97*. Error bars are SD. *** t-test, P<0.001. (**I**) *Sirt2* mutant spermatocytes display centriolar extensions positive for Ana1. However, the centriole barrel itself is not significantly elongated. Scale bar is 1µm. (**J**) Quantitation of defects in (**I**). Error bars are SD. *** t-test, P<0.001. (**K**) Model for Cep97 function. Cep97 is stably recruited to mature, full-length centrioles. Cep97 acts to suppress further centriolar microtubule dynamics in part via interaction with the microtubule acetylation machinery.

The phenotypic similarities with Cep97 prompted us to examine centrioles and basal bodies in acetylation defective animals. In wing discs, *Sirt2* mutants displayed severely truncated centrioles to an extent similar to *Cep97*. Interestingly, so did mutants of *Hdac6* and the acetylase *Atat1*. Centriole lengths were also more variable, again as with *Cep97* (Fig. 4D, E). One notable difference was the absence of extensions of individual centriolar microtubules found with *Cep97* and *CP110* (Fig. 4F). Combined loss of the acetylation machinery and Cep97 did not result in any further shortening of centrioles. However, the centriolar microtubule extensions found in *Cep97* single mutants were largely absent in *Cep97;Atat1* double mutants, suggesting they are destabilized (Fig. 4D-F). In spermatogenesis, *Atat1*, *Sirt2* and *Hdac6* mutant basal bodies did not display the over-elongation phenotype observed with *Cep97*, although lengths were more variable than in controls (Fig. 4G, H). *Sirt2* mutants did, however, display apparent basal body extensions positive for the centriolar marker Ana1 (Fig. 4I, J).

Thus, the phenotype of Cep97 and components of the acetylation/deacetylation machinery is remarkably similar though not identical, suggesting a shared function in regulating centriolar microtubule stability. This prompted us to examine the acetylation state of centriolar microtubules in the various conditions. Available reagents for acetylated tubulin show considerable non-specific background in the fly. However, basal body signal in sperm was found to be clearly dependent on Atat1, the major though not the only microtubule acetylase in the fly (Fig. 4G). Basal body acetylated tubulin signal was similarly reduced in *Cep97* mutants, as well as interestingly in mutants of *Sirt2* and *Hdac6* (Fig. 4G). Thus, the centriolar/ciliary phenotypes in *Cep97* mutants may at least in part be explained by disruption of centriolar microtubule acetylation. It may at first sight seem paradoxical that loss of the acetyltransferase Atat1 yields largely the same phenotype as loss of the deacetylases Hdac6 and Sirt2. However, this is not entirely unprecedented with both hyper-acetylation [48] and -glutamylation [49], another normally stabilizing post-translational modification, being reported to destabilize microtubules, the implication being that a large imbalance in modification is as destabilizing as complete loss of the modification.

### A revised model of Cep97 function

In summary, our results indicate that Cep97 forms part of a capping structure that is stably recruited to fully elongated, mature centrioles. Removal of Cep97 need not occur to allow initiation of the ciliary axoneme. Rather, Cep97 is required for proper basal body function in ciliogenesis. Consistent with its late loading onto centrioles, loss of Cep97 does not affect procentriole assembly up to and including formation of the CPAP/Sas-4-containing centriole wall. However, failure to assemble the Cep97 cap structure leaves centriolar microtubules exposed, which can result in abnormal extension or shrinkage of centrioles depending on cytoplasmic context. Instead of counteracting the activity of centriole elongation factors such as CPAP, CEP120 or SPICE1, Cep97 therefore acts to limit microtubule dynamics once the proper length has been reached. Our work provides novel mechanistic insight into how this may occur. A hallmark feature of centrioles is their remarkably constant size and stability, with no detectable turnover of tubulin subunits subsequent to their incorporation [50], enabling individual centrioles to be traced throughout many cell divisions [51] and indeed over the lifetime of an animal [52]. The extensive post-translational modification of centriolar microtubules, including by acetylation, is likely to be key for this stability, as memorably demonstrated by the dissolution of centrioles following microinjection of antibodies against poly-glutamylated tubulin [53]. Here, we identify the microtubule deacetylase Sirt2 as a Cep97 interactor. We further show that loss of Cep97 results in a loss of centriolar microtubule acetylation. Remarkably, perturbation of components of the microtubule acetylation machinery (Sirt2, as well as another deacetylase Hdac6 and the acetyltransferase Atat1) largely phenocopies the effect of loss of Cep97. All three components have been localized to stable microtubules, including those at centrioles [42, 54, 55]. The most simple explanation, then, is that Cep97 regulates this machinery at centrioles to stabilize centriolar microtubules and confer on them their remarkable lack of dynamics, so different from that of other cytoplasmic microtubules (Fig. 4K). Finally, it bears remarking that the effects of Cep97 perturbation are much more severe than for CP110 [17]. Cep97 is also more widely conserved across eukaryotes than CP110, which is found exclusively in metazoans [56]. Cep97 is therefore clearly much more than just a CP110 recruitment factor. Yet it is CP110 that has attracted the most attention. The work presented here should serve as a foundation for future studies examining Cep97 function also in other experimental models.

## EXPERIMENTAL PROCEDURES

### *Drosophila melanogaster* stocks and husbandry

All strains used are listed in the Key Resource Table. *w^1118^* flies were used as wild-type controls. Sas-4-GFP [57], *CP110Δ*, and CP110-GFP [17] strains have been described previously. Mutants in *Atat1*, *Hdac6* and *Sirt2* were obtained from the Bloomington *Drosophila* Stock Center (BDSC). The *Cep97* null mutant was created by ends-out or replacement gene targeting [34] using the pGX-attP vector [58]. Briefly, 5kb fragments upstream of the start codon and downstream of the stop codon of the Cep97 gene were cloned on either side of the white gene in the pGX-attP plasmid. This plasmid was then injected into early embryos and flies selected for red eyes. I-SceI endonuclease was then activated by crossing to SceI carrying flies and offspring with red eyes selected. Cep97 deletion was confirmed by Southern, PCR and sequencing. Endogenous promoter Cep97-GFP was created by recombineering. A GFP tag was inserted at the C-terminus of Cep97 into BAC RP98-06H02 via bacterial Red/ET recombination (Gene Bridges). Cep97-GFP together with 3.5kb of upstream and 4.2kb of downstream regulatory sequences was transferred to the injection vector pCM43 (gift of Barry Dickson), again by homologous recombination, and inserted into the landing site attP2 as previously described [59]. All Cep97 localization studies were performed in a *Cep97*-deletion mutant background. GFP-Cep290 was constructed by CRISPR-mediated genome editing. A guide-RNA to the N-terminus of Cep290 and rescue construct with 1kb flanking arms was injected into flies and positive clones identified by PCR and sequencing.

### Cell culture

Immortalized human retinal pigment epithelial cells stably expressing telomerase reverse transcriptase (hTERT-RPE1/RPE cells) were cultured in Dulbecco’s modified Eagle’s medium (DMEM) containing 10% FCS, penicillin (100 units/ml), and streptomycin (0.1mg/ml). Cells were grown at 37°C in 5% CO2 in air and passaged every 2-3 days. For both immunofluorescence staining and expansion microscopy cells were seeded on glass coverslips. To induce ciliation cells were serum starved for 24h prior to fixation. HEK293T cells were cultured as above. Full length Cep97 was amplified from HEK293T cDNA and cloned into the vector p3xFLAG-CMV-10 (Sigma). SIRT2 was transferred from the Gateway donor vector pDONR221 [40] into the destination vector pDEST 2xHA (gift of Joanna Loizou). Transient transfections of HEK293T cells were performed with polyethylenimine (Polysciences).

### *Drosophila* behavioral assays

#### Larval touch assay

This assay was performed as previously described [35]. In brief, third-instar larvae moving forward were touched with the tip of an eyelash across one side of the thoracic segments and the response measured on a scale of 0-4 (0 no response, 1 hesitation, 2 turn, 3 reversal, 4 complete reversal). This was repeated 4 times for each animal and the total used to calculate a reactive score from 0-16. Experiment was repeated 3 times with >10 larvae per condition.

#### Climbing assay

To examine fly coordination, 12 two-day-old adult males were collected in 16cm graduated, flat bottom tubes. Flies were allowed to recover from the anesthetic and left in the tube for 15min to acclimatize, then banged to the bottom of the tube and filmed climbing back upwards. Videos were analyzed to establish the time at which 10 flies had crossed the half-way (8cm) mark. Banging was repeated four times and the average time recorded. Experiment was repeated 3 times with 12 males per condition.

#### Male fertility assay

To test the fertility of male flies, single males were crossed with four virgin control females in standard culture vials. Flies were mated for 1h. Males were then removed and females left in the vial for 24h. Numbers of offspring were counted prior to hatching. Experiment was repeated 3 times with >10 males per condition.

### Antibodies for immunofluorescence and immunoblotting

Rabbit polyclonal antibodies against the N-terminus (amino acids 1-328) and C-terminus (631-806) of *Drosophila* Cep97 were raised using an MBP fusion as an antigen and purified over a GST fusion as described [60]. The following primary antibodies were used for immunofluorescence in *Drosophila*: GFP-booster-Atto488 (ChromoTek) and mouse anti-acetylated tubulin (6-11B-1; Sigma-Aldrich) at 1:200, rabbit anti-Cep97-N-ter and anti-Cep97-C-ter, rabbit anti-Asl and rabbit anti-Ana1 [61], rabbit anti-Sas4 [62] and mouse anti-NompC [63], all at 1:500; mouse anti-*γ*-tubulin (GTU88; Sigma-Aldrich) and mouse anti–α-tubulin (DM1α; Sigma-Aldrich), both 1:1000. The following secondary antibodies were used: Alexa Fluor 488, 568 and 647 (Invitrogen), all at 1:1000. To stain actin Phalloidin (Alexa 568) was used at 1:500. DNA was labeled with Hoechst 33342 (Invitrogen). For STED microscopy we used GFP-booster-Atto594 at 1:200 and STAR RED-conjugated secondary antibodies at 1:500. For immunofluorescence in RPE cells the following antibodies were used: mouse anti-acetylated tubulin (C3B9 [64]; gift of Vladimir Varga) at 1:100, rabbit anti-hsCep97 (Proteintech) at 1:150 and rabbit anti-hsCP110 (Proteintech) at 1:500. Anti-mouse Alexa Fluor 488 and anti-rabbit Alexa Fluor 555 secondary antibodies (Invitrogen) were used at 1:1000. For immunoblotting, rabbit anti-Cep97-C-ter antibodies (this study) were used at 1:500, mouse anti-HA at 1:1000, mouse anti–α-tubulin (DM1α; Sigma-Aldrich) at 1:1500 and mouse anti-FLAG-HRP at 1:10000. HRP-coupled secondary antibodies were used at 1:7500.

### Immunoprecipitation

HEK293T cells were co-transfected with 4µg of HA-SIRT2 and 4µg of either FLAG empty vector or FLAG-CEP97. Cells were harvested 48h after transfection and lysed in lysis buffer (50mM Tris-Cl pH 8, 150mM NaCl, 1% Triton, 1x Roche Complete Mini Protease Inhibitor Cocktail, 1mM PMSF, 5μM trichostatin A, 20mM nicotinamide, 50units/mL benzonase and 1mM DTT) for 1h at 4°C. 10% of the cleared lysate was kept as input and the rest incubated for 2h on a rotating wheel at 4°C with anti-FLAG M2 magnetic beads (Sigma). Beads were subsequently washed three times with lysis buffer and immunoprecipitated proteins eluted with FLAG peptide.

### Southern and Western blotting

#### Southern

To identify deletion mutants, 20µg of genomic DNA were digested using SmaI for 4 hours. Bands were separated on a 0.7% polyacrylamide gel and transferred to nylon membrane, incubated with a P32-labelled probe and washed. Finally, a phosphor-storage screen was exposed to the membrane and developed using a phosphorimager.

#### Western

To examine Cep97 protein levels in *Cep97* mutants, dechorionated embryos were ground with a pestle and mortar and boiled for 10min in loading buffer. Embryo lysates were separated on an 8% polyacrylamide gel and transferred to a nitrocellulose membrane. Blots were probed with anti-Cep97 and anti-tubulin antibodies overnight at 4°C, followed by 2h incubation with HRP-conjugated followed by 2h incubation with HRP-conjugated secondary antibodies and bands visualized by ECL on a Bio-Rad ChemiDoc Touch Imaging System. For Western blot analysis of FLAG-hsCep97 immunoprecipitates, 5% of the input and 30% of the eluate were loaded for each sample on a 10% gel and transferred onto nitrocellulose membrane. Blots were probed with anti-FLAG-HRP and anti-HA antibodies overnight at 4°C, followed by 1h incubation with anti-mouse-HRP secondary antibody in the case of anti-HA antibody. Bands were visualized by ECL.

### PCR and RT-PCR

PCR was performed on fly lysates according to standard protocols. For RT-PCR, mRNA was prepared from 10 dissected brains or 10 testes using the RNeasy kit (Qiagen). cDNA was then generated using the QuantiTect Reverse Transcription kit (Qiagen).

### Immunofluorescence staining of *Drosophila* tissues

Cep97-GFP was detected with anti-GFP. All antibodies were used at dilutions detailed above.

#### Embryos

Embryos were dechorionated and formaldehyde fixed as previously described [61].

#### Larval brains

Brains were dissected from 3^rd^ instar larvae and fixed as previously described [62].

#### Wing discs

Wing discs were dissected from 3^rd^ instar larvae and fixed in 4% formaldehyde in PBS 0.1% triton (PBST) for 20min. Samples were then incubated for 15min in PBST, washed twice with PBS, blocked in 5% BSA in PBST for 30min before incubating with primary antibodies as above.

#### Leg chordotonal organs and testes

Legs chordotonal organs were dissected from 36h old male pupae and testes from 72h old male pupae. Fixation and staining were done as previously described for testis [65]. For actin staining, testes were fixed for 25min in 4% formaldehyde in PBS 0.1% triton. Samples were then incubated for 10min in PBST, washed twice with PBS, blocked in 5% BSA in PBST for 30min before incubating with phalloidin and primary antibodies as above.

### Immunofluorescence staining of RPE1 cells

#### Immunofluorescence staining of non-expanded cells

Coverslips were fixed with 4% formaldehyde in PBS for 10min at room temperature and briefly washed with PBS. Fixed cells were permeabilized with 0.5% Triton X-100 in PBS for 5min and then washed 3x in PBS. Cells were stained with primary antibodies diluted in 2% BSA in PBS for 1h in a humid dark chamber. After 3x washes with PBS, coverslips were stained with secondary antibodies diluted in 2% BSA in PBS for 1h. Finally, coverslips were washed in PBS 2x for 5min and 1x with ddH2O. After air-drying, coverslips were mounted in DAPI-containing ProLong Gold anti-fade mounting medium (Invitrogen).

#### Expansion microscopy

Protocol is based on [66]. Coverslips with cells were fixed with 4% formaldehyde/4% acrylamide in PBS overnight and then washed 2x with PBS. The gelation was performed by incubating coverslips face down with 45μl of monomer solution (19% (wt/wt) sodium acrylate, 10% (wt/wt) acrylamide, 0.1% (wt/wt) N,Ń-methylenbisacrylamide in PBS supplemented with 0.5% TEMED and 0.5% APS, prepared as described in [66]) in a pre-cooled humid chamber. After 1min on ice, chamber was incubated at 37°C in the dark for 30min. Samples in gel were denatured in denaturation buffer (200mM SDS, 200mM NaCl, 50mM Tris in ultrapure water) at 95°C for 4h. Gels were expanded in ddH2O for 1h until they reached full expansion factor (4.2x) and then cut into 1×1cm pieces. Pieces of gel were incubated with primary antibodies diluted in 2% BSA in PBS overnight at RT. After staining, shrunk pieces of gel were incubated in ddH2O for 1h, during which time they re-expanded. After reaching their original expanded factor, pieces of gel were incubated with secondary antibodies diluted in 2% BSA in PBS for 3h at RT. Last expansion in ddH2O with exchange every 20min was for 1h until pieces of gel reached full size. Samples were imaged in 35mm glass bottom dishes (CellVis, USA) precoated with poly-L-lysine. During imaging, gels were covered with ddH2O to prevent shrinking.

### Fixed and live cell imaging

#### Standard confocal microscopy

Immunofluorescence samples were analyzed on a Zeiss LSM710 scanning confocal microscope equipped with an Airyscan unit. Stacks of 0.75µm slices were acquired with a 63x 1.4NA Plan Apochromat lens using single channel mode to avoid cross-illumination. For centriole length measurements in testes, 4x electronic magnification was used. Airyscan images were acquired for selected centrioles using 0.25µm slices and 10x electronic magnification. Maximum intensity projections of Z-stacks prepared in ImageJ were used for image analysis and panel preparation.

#### STED microscopy

STED images were acquired on an Abberior STEDYCON unit on a Zeiss Axio Imager A2 microscope. Image stacks of 0.1µm slices were acquired with a 100x 1.46NA alpha Plan-Apochromat lens and using 561/775nm and 640/775nm excitation/depletion combinations. Maximum intensity projections of Z-stacks prepared in ImageJ were used for image analysis and panel preparation.

#### Live cell imaging and FRAP analysis

Dechorionated embryos were covered with Voltalef and examined on a Yokogawa CSU X1 spinning disk confocal mounted on a Zeiss Axio Observer Z1 inverted microscope equipped with a 63x 1.4NA Plan Apochromat lens, 120mW 405nm and 100mW 488nm solid-state lasers, 2D-VisiFRAP Galvo FRAP module and Photometrics CoolSNAP-HQ2 cooled CCD camera and controlled by VisiView software (Visitron Systems). Z-stacks of 0.75µm were acquired every 15 seconds. Photobleaching was performed using the galvanometer point scanner to target a region encompassing multiple centrosomes with the 405nm laser at 120mW power. Image stacks were imported into ImageJ for post-acquisition processing.

#### Image acquisition RPE cells

Expanded cells were imaged by confocal microscopy on a Leica TCS SP8 scanning confocal microscope using a 63x 1.4NA oil objective with closed pinhole to 0.5 AU. Cilia were acquired in Z-stacks at 0.1μm stack size with pixel size 36nm. Images were computationally deconvolved using Huygens Professional software (Scientific Volume Imaging) prior to for image analysis and panel preparation.

### Transmission electron microscopy

#### Chordotonal organs

Legs from 36h old pupae were cut off with microscissors and fixed using a mixture of 2% glutaraldehyde and 2% paraformaldehyde in 0.1 mol/l sodium phosphate buffer, pH 7.2 for 2h in a desiccator at room temperature and then overnight on a rotator at 4°C. Legs were then rinsed with sodium phosphate buffer, post-fixed in 2% osmium tetroxide in buffer on ice, dehydrated in a graded series of acetone on ice and embedded in Agar 100 resin. 70nm sections were cut and post-stained with 2% uranyl acetate and Reynolds lead citrate. Sections were examined with a Morgagni 268D microscope (FEI, Eindhoven, The Netherlands) operated at 80kV. Images were acquired using an 11 megapixel Morada CCD camera (Olympus-SIS).

#### Testes/Wing discs

Late pupal testes and wing discs from third instar larvae were dissected in PBS and fixed using 2.5% glutaraldehyde in 0.1mol/l sodium phosphate buffer, pH7.2 for 1h at room temperature. Samples were then rinsed with sodium phosphate buffer, post-fixed in 2% osmium tetroxide in dH_2_O on ice, dehydrated in a graded series of acetone and embedded in Agar 100 resin. 70nm sections were then cut and processed as above.

### Image analysis

#### Centriole length

Centriole length was measured on maximum-intensity z-projections of testes stained with Ana1 or Asl. At the spermatocyte stage, perfectly oriented centriole pairs (V-shape clearly visible) were selected and only the mother centriole measured in Image J, with beginning and end of the centriole defined as where the signal reached 25% of the maximum intensity. >25 centrioles measured per condition.

#### Photobleaching experiments

Maximum projected stacks were opened and bleach-corrected in ImageJ (simple ratio matrix). Bleached and unbleached centrioles were tracked manually. A circle with a diameter of 7 pixels was used to encompass the centriole and average intensities measured over time for each centriole. Background subtraction was done by measuring the average signal in a 20 pixel circle of nearby cytoplasm. Two bleached centrioles and four unbleached control centrioles were measured per embryo and the signal averaged for each condition. Plots are of 3-point moving average to smooth out signal intensity variation due to centriole movement in z. Time of initial recovery was defined as the time at which centriolar signal initially became detectable again after photobleaching. Time of completion of recruitment was defined as the time at which plateau levels (∼50% of original signal) were first reached for three successive frames.

#### Centriolar signal quantitation

To measure the lateral distribution of Cep97 relative to other centriolar markers in *Drosophila* spermatocytes and expanded RPE cells, linescans were performed in Image J on maximum-intensity z-projections of perfectly oriented centrioles, that is ones captured lying in or perpendicular to the plane of imaging. Where proteins were located peripherally near the wall of the centriole barrel, the distance between the two peaks of fluorescence intensity was measured for each centriole. For proteins located towards the central lumen of the centriole, distribution was measured as the full width at half maximum. For quantitations in RPE cells, values were divided by the expansion factor of 4.2 to obtain original non-expanded dimensions.

### Quantification of centriole and axonemal defects in electron micrographs

#### Wing discs

Centrioles of three wing disc were analyzed per condition. Images were acquired at 36000x for >15 centrioles per wing disc and length and diameter measured using ImageJ.

#### Leg chordotonal organs

Leg chordotonal organs from three different animals were analyzed per condition. Cross sections or longitudinal sections of leg chordotonal cilia were analyzed and scored for presence or absence or other defects at 72000x and 36000x magnification.

#### Testes

Three testes were analyzed per condition. To detect gross defects in cell division, cross sections of whole cysts were analyzed at low magnification (4000x-8000x). Sperm axoneme abnormalities were analyzed at 36000x. Missing or defective axonemes per cyst (normal 64 axonemes) were scored for >20 cysts per condition.

### Statistical Analysis

All error bars are standard deviation. To compare samples in a specific experiment, we conducted unpaired t-tests using GraphPad Prism. *, **, *** represent P-Values of <0.05, 0.01 and 0.001, respectively. Tests are comparing indicated condition to control unless otherwise specified.

## KEY RESOURCES TABLE

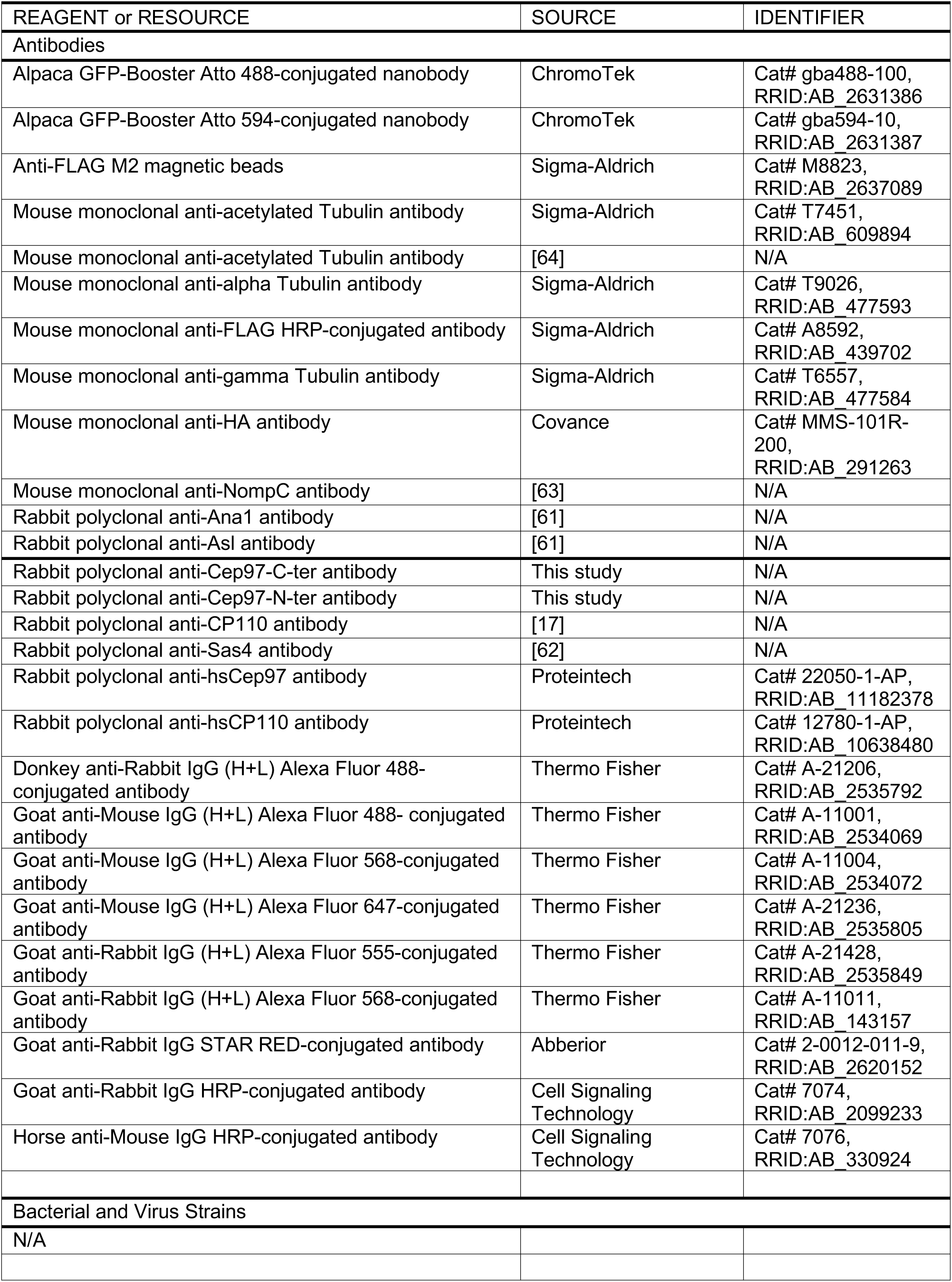
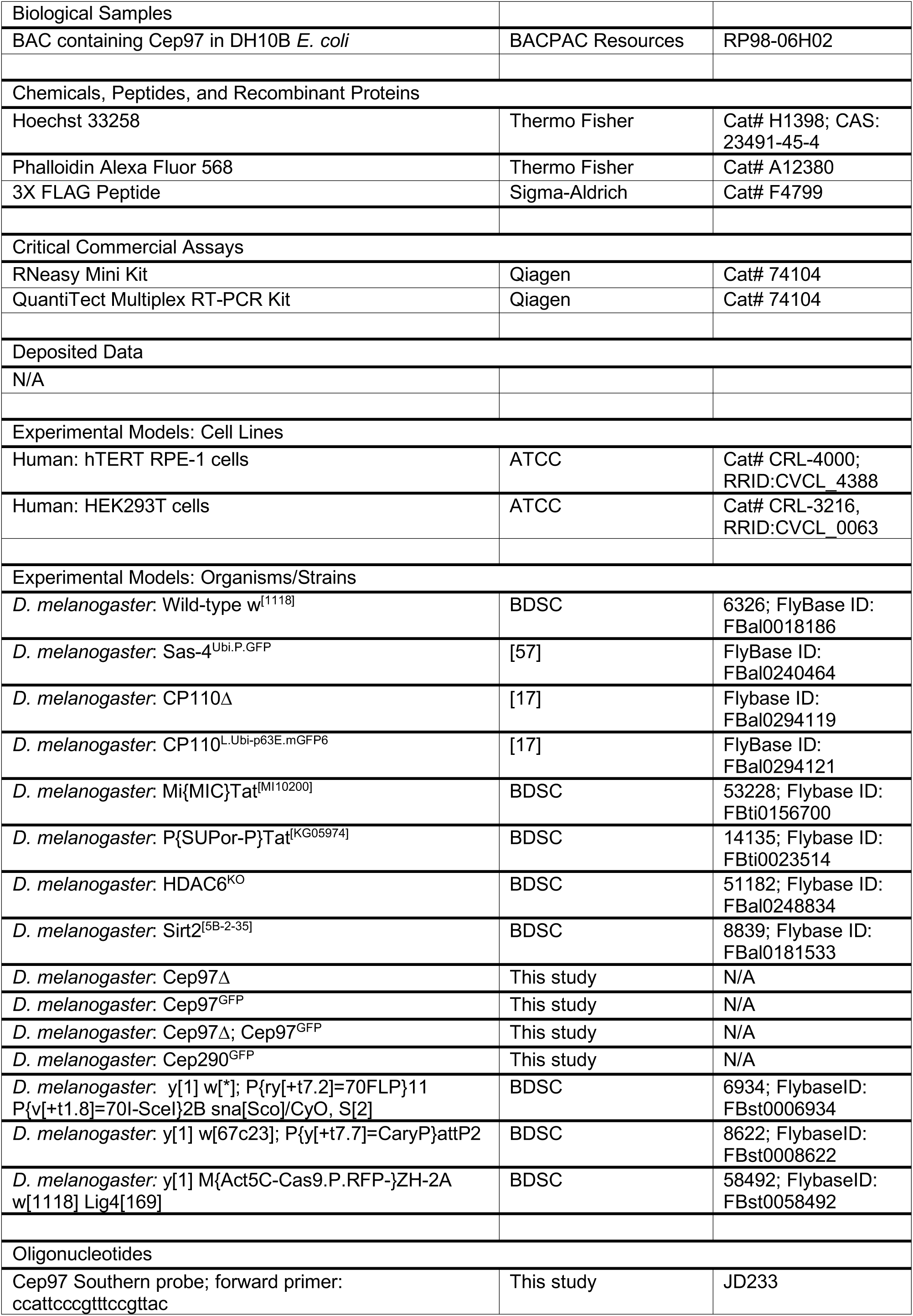
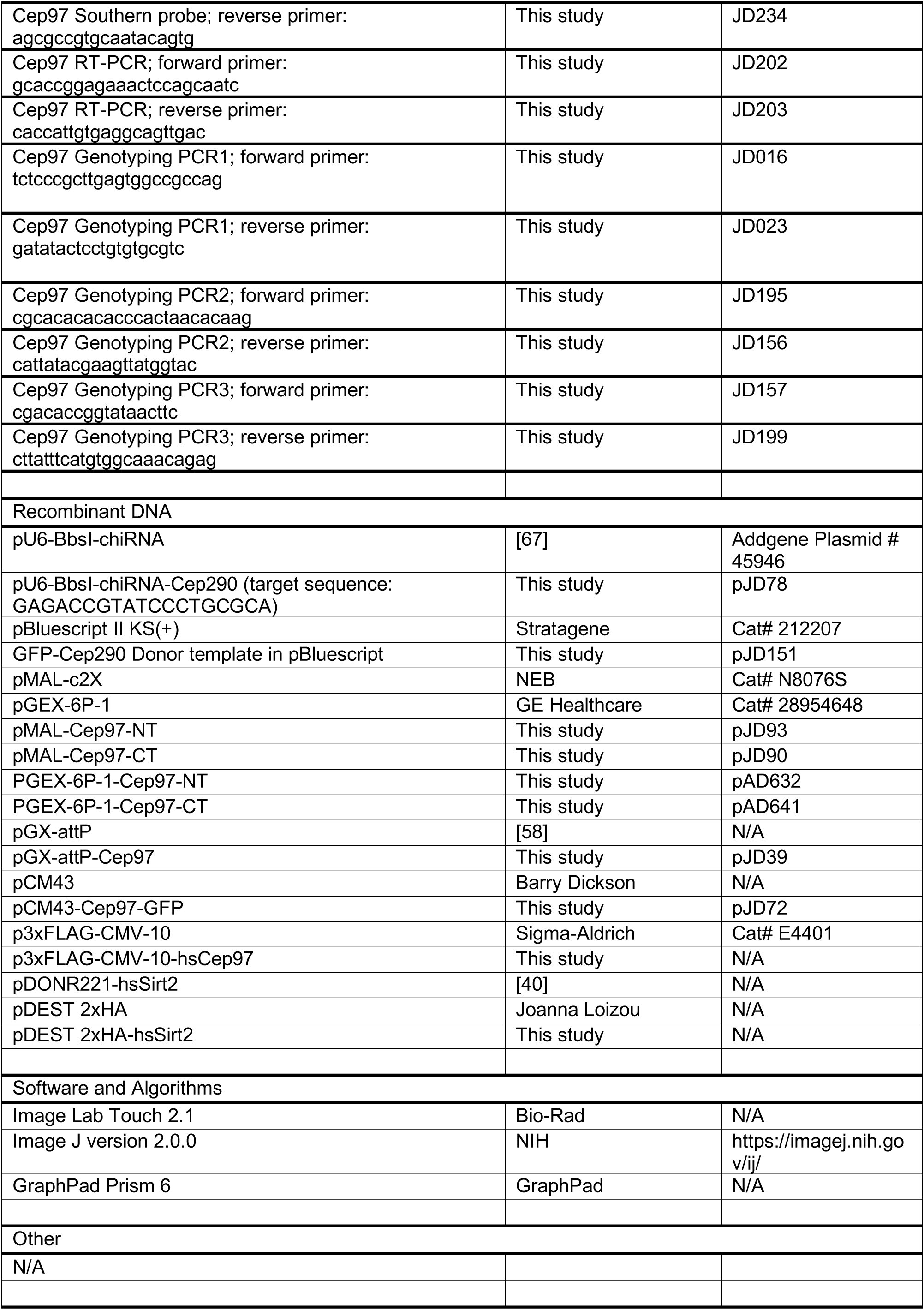

## ACKNOWLEDGEMENTS

We thank members of the Dammermann and Slade labs for discussions; Balazs Erdi, Peter Duchek and Joseph Gokcezade of the IMBA fly house and members of the Dickson and Knoblich labs for help with the *Drosophila* experiments; Nicole Fellner of the VBCF Electron Microscopy facility for help with preparing samples for EM; the Bloomington *Drosophila* Stock Center, Joe Howard, Joanna Loizou, Jordan Raff, David Stanek, Vladimir Varga and the Vienna *Drosophila* Resource Center (VDRC) for strains and reagents; and Josef Gotzmann and Thomas Peterbauer of the MFPL BioOptics facility for technical assistance. This work was supported by grants Y597-B20 from the Austrian Science Fund (FWF) to A.D. and 17-20613Y from the Czech Science Foundation to M.S.-C. and M.H., as well as a Lisa-Meitner Fellowship of the FWF (M1293-B09) and a VIPS post-doctoral fellowship of the University of Vienna to J.D.

## AUTHOR CONTRIBUTIONS

J.D. conception and design, acquisition of data, analysis and interpretation of data, and drafting or revising the article; M.S.-C. and M.H. contributed unpublished essential data or reagents (RPE cell expansion microscopy); D.S. contributed unpublished essential data or reagents (Sirt2/Cep97 interaction in human cells); A.D. conception and design, analysis and interpretation of data, and drafting or revising the article.

## SUPPLEMENTAL FIGURE LEGENDS

**Figure S1:**
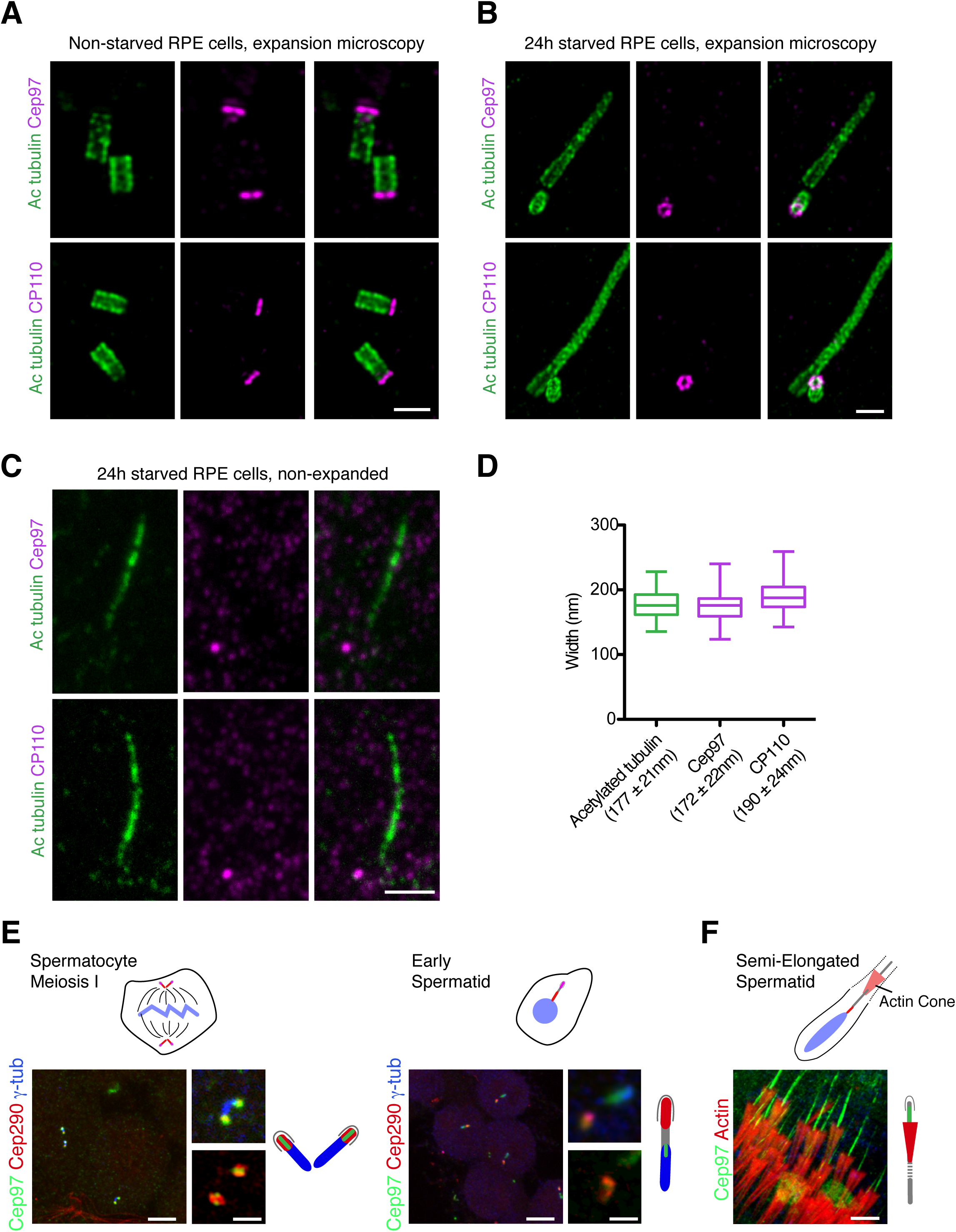
Further characterization of Cep97 localization. (**A-D**) Localization of CEP97 and CP110 in human RPE cells as assessed by expansion microscopy. 4.2x expanded RPE cells without starvation (**A**) or after 24h of serum starvation to induce ciliation (**B**). In cycling cells, CEP97 and CP110 co-localize with the centriolar wall marker acetylated tubulin at the distal ends of both mother and daughter centrioles. In starved cells, both proteins consistently localize to the distal end of daughter centriole only. Scale bars are 2μm. (**C**) Non-expanded serum-starved RPE cells stained with Cep97/CP110 and acetylated tubulin as above. Staining pattern is similar to in expanded RPE cells, but precise localization cannot be determined. Scale bar is 2μm. (**D**) Measurement of the lateral distribution of Cep97 and CP110 relative to the centriolar wall marker acetylated tubulin assessed on images as in (**A**). Values adjusted for expansion factor of 4.2. In contrast to in *Drosophila* (see Fig. 1B), Cep97 and CP110 localize close to the centriolar microtubule wall. Box and whisker plot with whiskers representing minimum and maximum values. N=49 Cep97, 46 CP110, 87 acetylated tubulin. (**E**) Cep97 localization during axoneme elongation in *Drosophila* spermatogenesis. Immunofluorescence micrographs of spermatocytes at meiosis I and early spermatids stained with Cep97 and the transition marker Cep290. While Cep290 moves with the growing axoneme tip [29], Cep97 largely remains at the distal end of centrioles marked with *γ*-tubulin. 2-color insets acquired in airyscan mode. Scale bars are 5µm, 1µm insets. (**F**) Cep97 prominently localizes to the axoneme ahead of the actin cones stripping cytoplasm from elongating sperm flagellum from base to tip [68]. Immunofluorescence micrographs stained with Cep97 and phalloidin to visualize actin. Scale bar is 5µm.

**Figure S2:**
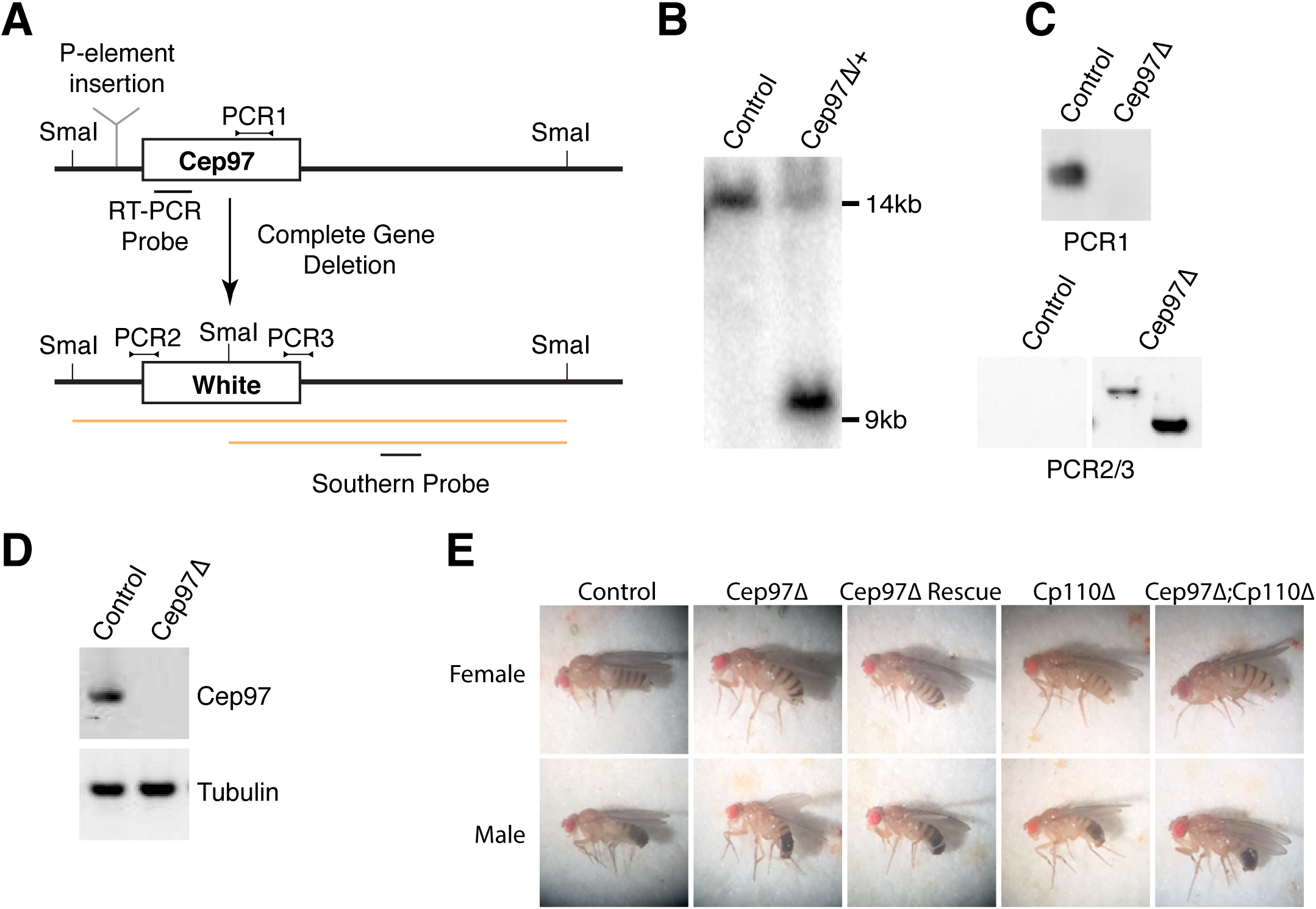
Generation of a *Cep97* null mutant. (**A**) Strategy used to generate *Cep97* deletion mutant. Previous work on Cep97 employed a transposable element insertion (*Cep97*^LL00167^, indicated) in the first intron of Cep97 ahead of the ATG [18]. Here, the entire coding sequence of Cep97 was replaced with the White marker gene. Schematic also indicates location of Southern, PCR and RT-PCR probes to confirm gene deletion/marker insertion. (**B**) Southern analysis shows expected band for successful gene replacement in original *Cep97* mutant heterozygote animals. (**C**) PCR confirms loss of Cep97 coding sequence and replacement by White marker. (**D**) RT-PCR reveals no residual expression of Cep97 in mutant animals. Tubulin used as loading control. (**E**) Appearance of Cep97/CP110 single and double mutant animals, as well as *Cep97* mutants rescued by GFP transgene. *Cep97* and *CP110* mutants display abnormal wing posture, a phenotype associated with defective mechanosensation.

**Figure S3:**
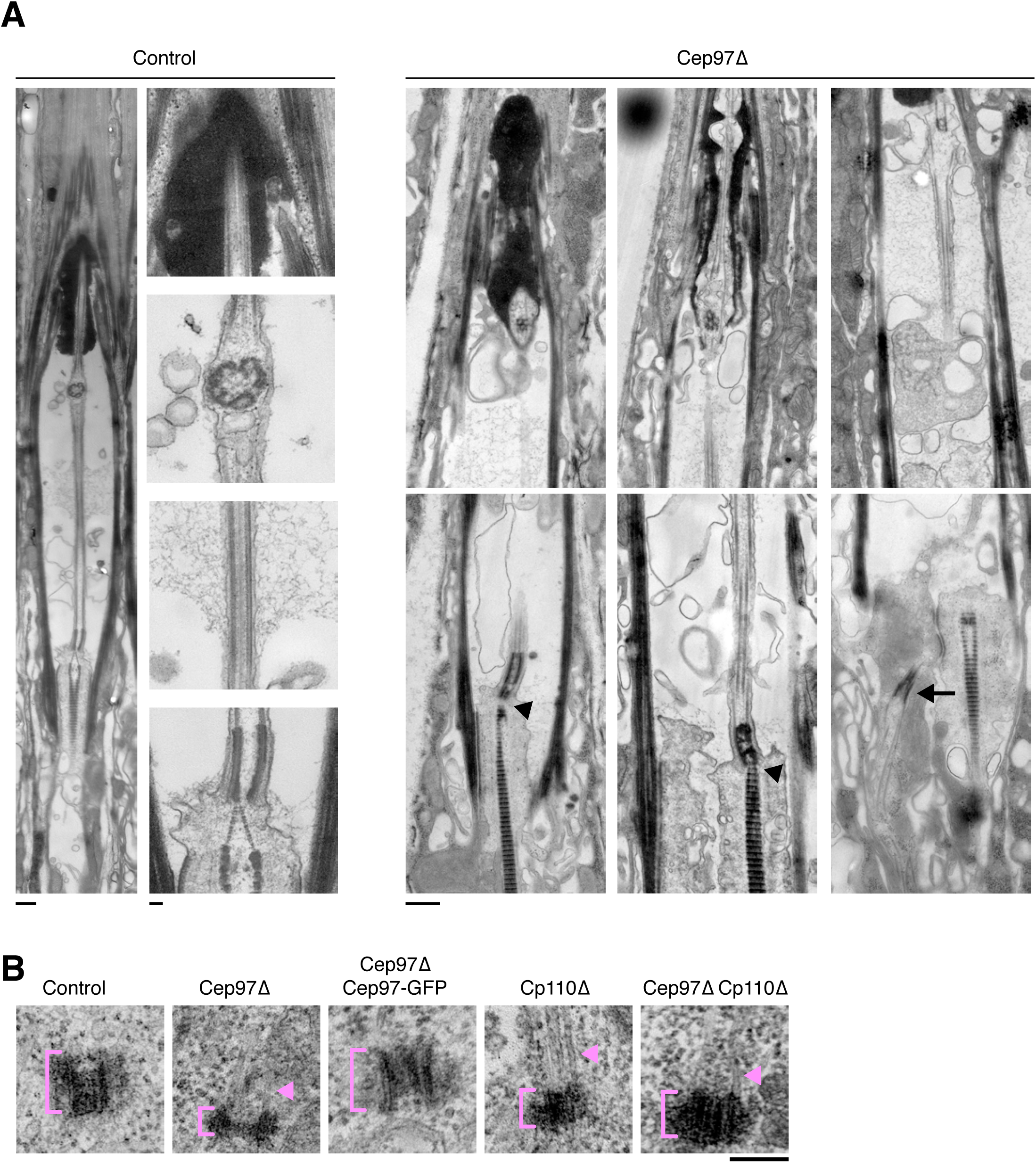
Centriole and ciliary ultrastructure of *Cep97* null mutants. (**A**) Transmission electron micrographs of scolopidia of wild-type and *Cep97* mutant leg chordotonal neurons. Longitudinal sections including expanded views of images shown in Fig. 2F to illustrate features of wild-type cilia (axoneme, ciliary dilation, basal body and daughter centriole decorated by rootlet) and range of defects in *Cep97* mutants. These include mispositioned basal bodies (arrow), missing daughter centriole and rootlets emanating directly from basal body (arrowheads). Distal features including ciliary dilation (when present) appear largely normal. Scale bars are 500nm, 100nm insets in control. (**B**) Transmission electron micrographs of centrioles in control, *Cep97* mutant, *Cep97Δ* Rescue, *CP110* mutant and *Cep97;CP110* double mutant wing discs. Note microtubule extensions in both Cep97 and CP110 mutants (arrowheads). See also Fig. 3C-F. Scale bar is 100nm.

**Figure S4:**
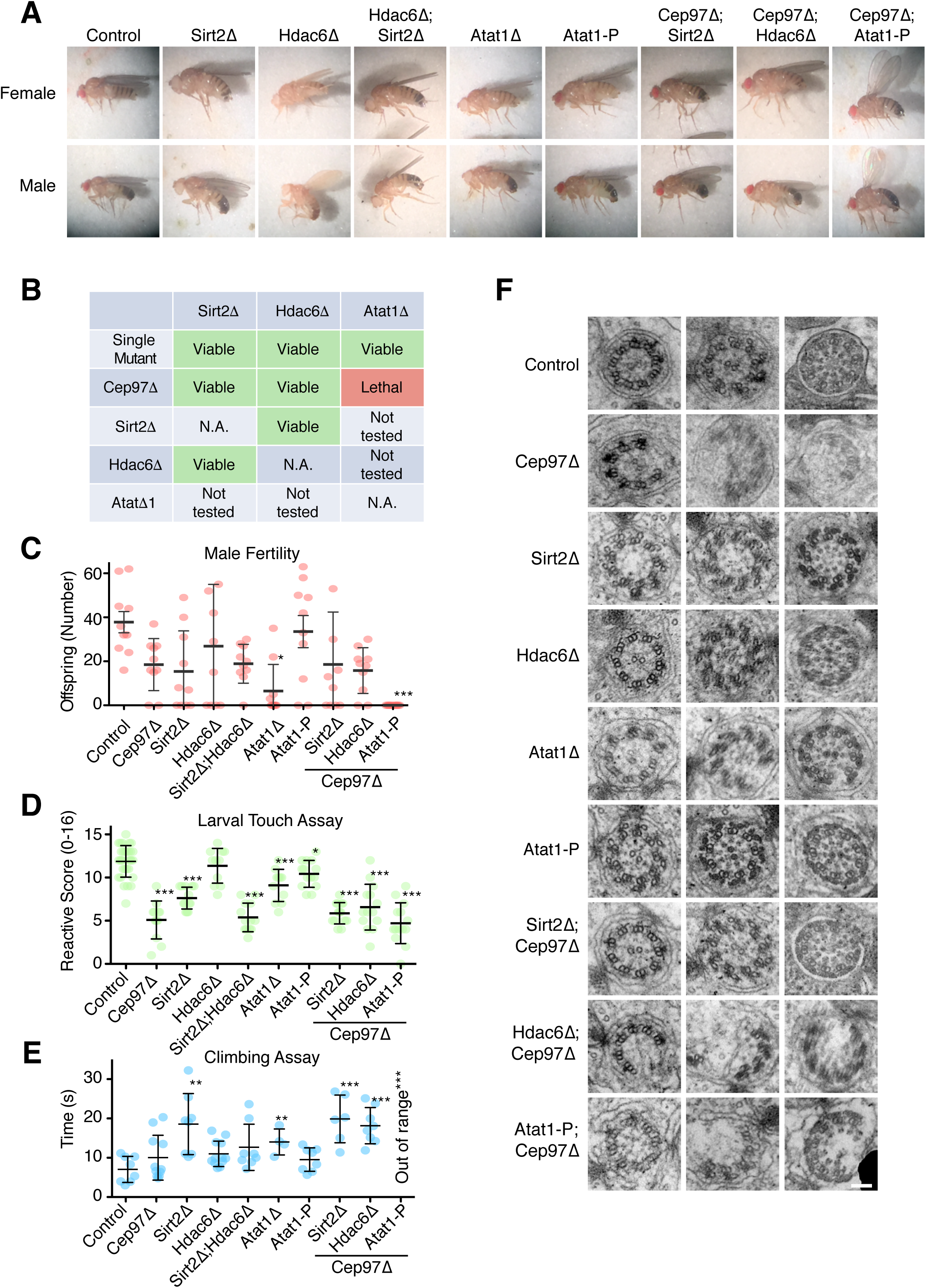
Further characterization of acetylation mutants. (**A**) Appearance of single and double mutants of *Sirt2*, *Hdac6* and *Atat1*. Mutants display abnormal wing posture, indicative of defective mechanosensation. Phenotype is most severe in *Cep97;Atat1* double mutants, consistent with results in other assays. (**B**) Viability of single and double mutants between *Cep97* and acetylation machinery. *Cep97* and *Atat1* null mutants are synthetic lethal. Behavioral assays were therefore performed with *Atat1-P* hypomorphic mutants in combination with *Cep97*. (**C**) Acetylation mutants with the exception of *Atat1-P* hypomorphic mutants display reduced male fertility, assessed as number of offspring per single male. Phenotype is strongly synergistic for *Atat1* and *Cep97*. Error bars are SD. * t-test, P<0.05, *** P<0.001. (**D, E**) Behavioral assays in larvae (touch assay, **D**) and adult flies (climbing/bang assay, **E**) show uncoordination in acetylation machinery mutants, albeit to different degrees. Phenotype strongly enhanced in *Cep97;Atat1* double mutants. Error bars are SD. * t-test, P<0.05, ** P<0.01, *** P<0.001. (**F**) TEM analysis of sperm axonemes in control and *Cep97*, *Sirt2*, *Hdac6* and *Atat1* mutant testes. Cross sectional views of *Sirt2*, *Hdac6* and *Atat1* mutants reveal occasional reveal fragmented axonemes, a phenotype strongly enhanced in *Cep97;Atat1* double mutants. Scale bar is 100nm. Defects quantitated in Fig. 4C.

